# A critical role of retinoic acid concentration for the induction of a fully human-like atrial phenotype in hiPSC-CM

**DOI:** 10.1101/2023.01.03.522611

**Authors:** Carl Schulz, Muhammed Sönmez, Julia Krause, Edzard Schwedhelm, Pan Bangfen, Dzenefa Alihodzic, Arne Hansen, Thomas Eschenhagen, Torsten Christ

**Author notes:** Address for correspondence Professor Dr. med. Thomas Eschenhagen, E-Mail or PD Dr. med. Torsten Christ, Institute of Experimental Pharmacology and Toxicology, University Medical Centre Hamburg-Eppendorf, Martinistraße 52, 20246 Hamburg, Telefon: 040-7410-52414. Stem Cell Biology, Novo Nordisk A/S, Måløv, Denmark (Julia Krause).

## Abstract

Retinoic acid (RA) induces an atrial phenotype in human induced pluripotent stem cells (hiPSC), but expression of atrium-selective currents such as the ultrarapid (I_Kur_) and acetylcholine-stimulated K^+^ current (I_K,ACh_) is variable and less than in adult human atrium. We suspected methodological issues and systematically investigated the concentration-dependency of RA. RA treatment increased I_Kur_ concentration-dependently from 1.1±0.54 pA/pF (0 RA) to 3.8±1.1, 5.8±2.5 and 12.2±4.3 at 0.01, 0.1 and 1 µM, respectively. Only 1 µM RA induced enough I_Kur_ to fully reproduce human atrial AP shape and a robust shortening of action potentials (AP) upon carbachol. We found that sterile filtration caused substantial loss of RA. We conclude that 1 µM RA appears necessary and sufficient to induce a full atrial AP shape in hiPSC-CM in EHT format. RA concentrations are prone to methodological issues and may profoundly impact success of atrial differentiation.

## Introduction

Human induced pluripotent stem cell-derived cardiomyocytes (hiPSC-CM) represent a model to study electrophysiological consequences of gene variants and mutations on a human background. Consequently, several models have been developed to investigate ventricular arrhythmias. However, atrial fibrillation (AF) is much more common than ventricular arrhythmias and cannot yet be studied sufficiently in hiPSC-CM. Atrial-like hiPSC-CMs (hiPSC-aCM) resembling the electrophysiological properties of the human atrium could be used to investigate mechanisms of AF in vitro. In addition, hiPSC-aCMs could give fundamental insight into pacing-induced electrical remodeling, a technique frequently used in animals that lead to the widely accepted concept of AF-induced remodeling (Wijffels et al., 1995). Thus, recently developed protocols for the differentiation of hiPSC-aCM are an important progress in modeling atrial electrophysiology in a human setting (Goldfracht et al., 2020; Laksman et al., 2017; Soepriatna et al., 2021).

Retinoic acid (RA) has been identified as a critical factor during atrial differentiation of hiPSC (Devalla et al., 2015; Lee et al., 2017). Indeed, inclusion of RA in standard differentiation protocols induced several electrophysiological parameters indicating an atrial, rather than ventricular phenotype of the resulting hiPSC-aCM. The presence of the G-protein-gated K^+^ channel (I_K,ACh_) is a hallmark of atrial and absent in ventricular cardiomyocytes (Heidbüchel et al., 1987) and therefore researchers have looked for shortening of APD_90_ upon activation of muscarinic receptors, but effect sizes were small and variable in two studies (Devalla et al., 2015; Lemme et al., 2018). Large mammals (dog and human) also possess large transient potassium currents consisting of the transient outward current (I_to_) and the atrial-selective, ultrarapidly activating potassium current (I_Kur_) (Burashnikov et al., 2004; Ravens and Wettwer, 2011). These currents dominate the early repolarization phase and lead to the typical spike and dome shape of the atrial action potential (AP). In notable contrast, hiPSC-aCM resulting from various RA-supplemented differentiation protocols exhibited a rather triangular AP shape without the prominent spike (Argenziano et al., 2018; Goldfracht et al., 2020; Gunawan et al., 2021; Honda et al., 2021; Pei et al., 2017), indicating incomplete atrial differentiation. In our own hands, the AP shape of engineered heart tissue (EHT) from hiPSC-aCM showed strong inter-investigator variability, indicating methodological issues during the differentiation process. Given the established key role of RA, we set out to prospectively investigate the concentration-dependent effects of RA on amplitudes of the atrial selective I_Kur_ in hiPSC-aCM. The contribution of I_Kur_ and I_K,ACh_ to repolarization was estimated from AP recordings which were measured in intact aEHT.

## Results

To study the concentration-dependency of RA on atrial specification we added RA at 0.01, 0.1 or 1 µM after mesodermal induction of hiPSC for 3 days (Lemme et al., 2018). Cells cultured in the absence of RA were used as controls. We followed a standard embryoid body-based spinner protocol for cardiac differentiation of hiPSC (Breckwoldt et al., 2017). After EB dissociation, EHTs were casted, cultured for 4 weeks before they were used for either AP recordings or patch clamping in dissociated hiPSC-CM.

### Concentration-dependent effects of RA on outward currents

In human hearts, outward currents are larger in atrial than in ventricular cardiomyocytes, partially due to expression of the atrial-selective I_Kur_ in atria (Amos et al., 1996). In order to quantify effects of RA on atrial differentiation, we measured outward currents in hiPSC-CM differentiated in the presence of different concentrations of RA and in controls (differentiated in the absence of RA). We separated I_Kur_ from total outward current by applying a low concentration of the non-selective potassium channel blocker 4-AP (50 µM). Experiments were finished by 5 mM 4-AP to block not only I_Kur_ but also I_to_.

We found transient outward currents in all individual hiPSC-CM (**Figure 1A, 1B**). In hiPSC-CM derived from a RA-free differentiation protocol, peak outward currents were not suppressed by 50 µM 4-AP, indicating absence of I_Kur_. Even the lowest concentration of RA (0.01 µM) was able to induce an outward current component sensitive to 50 µM 4-AP: I_Kur_. Cells treated with higher RA concentrations showed progressively more I_Kur_ (with 1 µM RA: 12.1±2.5, n=16) but also I_to_ (difference between the current in the presence of 50 µM 4-AP and the current in the presence of 5 mM 4-AP: 33.5±6.5 pA/pF, n=16) and in a 4-AP-insensitive outward current component.

**Figure 1:**
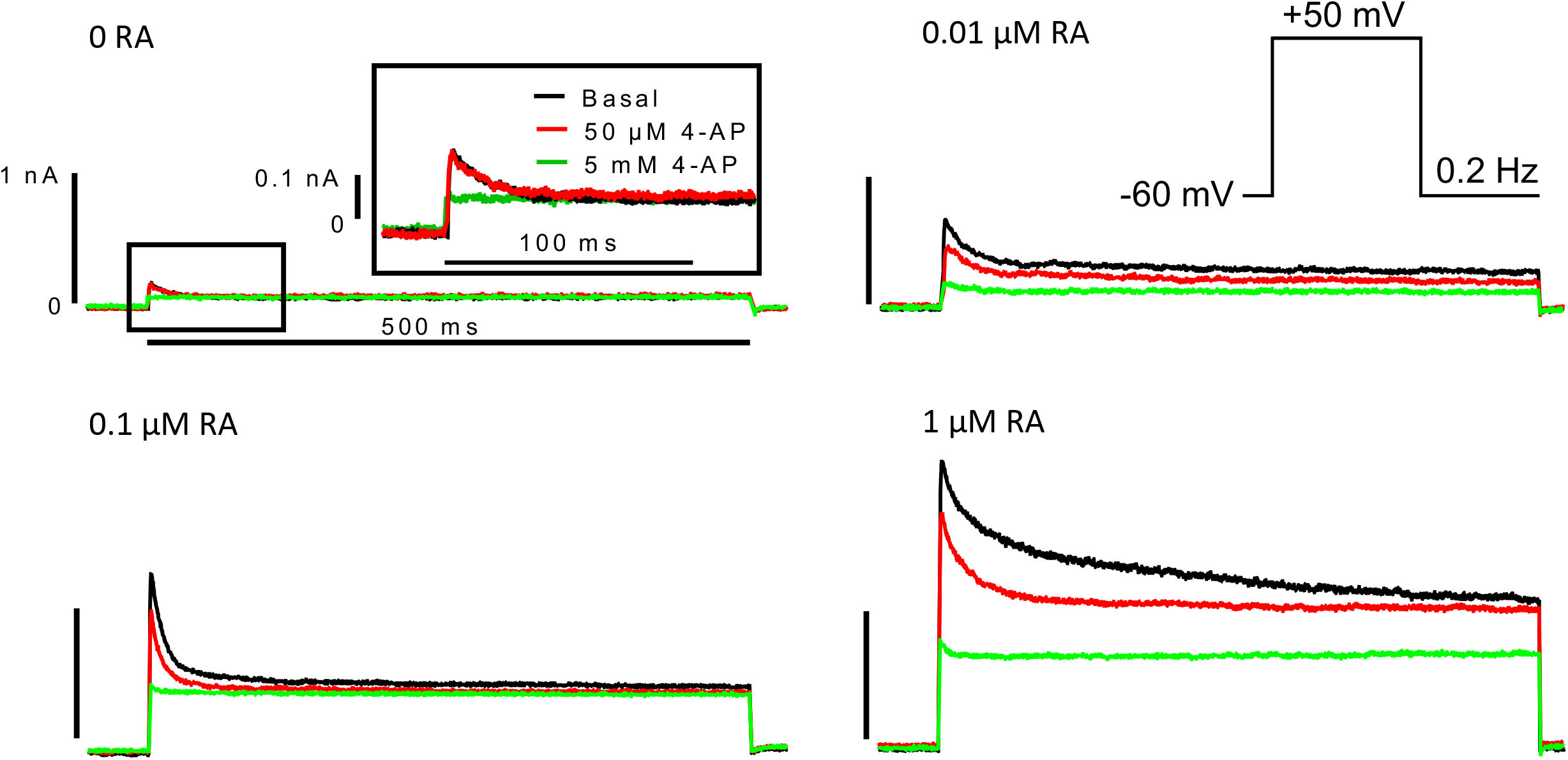
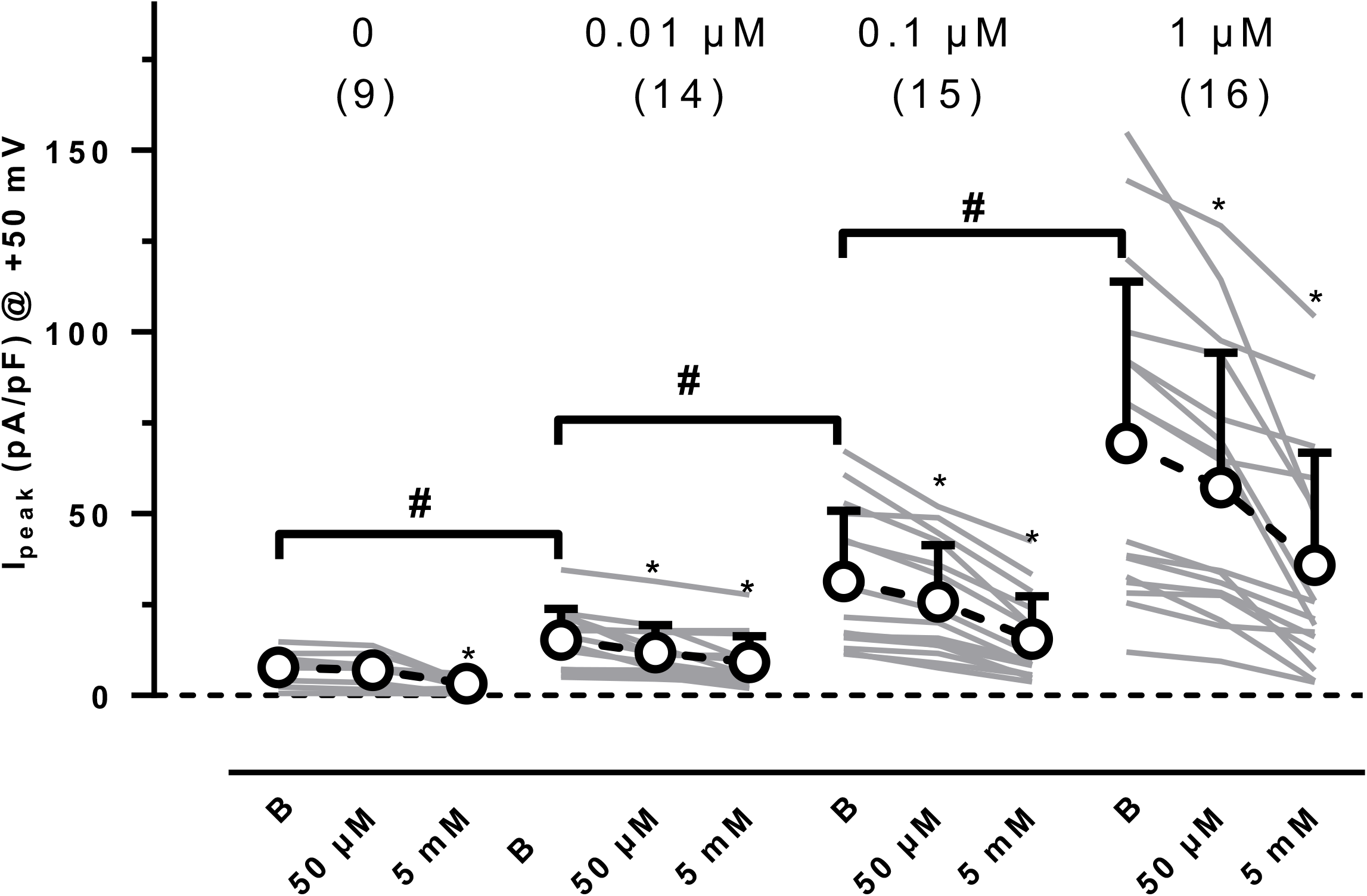
Concentration-dependency of RA on outward currents in hiPSC-CM. **A:** Original current traces recorded in individual hiPSC-CM differentiated in the absence (0 RA) or presence of different concentration of RA in in the absence (Basal) and in the presence of 50 µM 4-AP and 5 mM 4-AP. For 0 RA initial part of outward currents is given on an extended scale. Pulse protocol given as inset. **B:** Summary of the results for peak currents measured in the absence of 4-AP (basal, B) and in the presence of 50 µM and 5 mM 4-AP, recorded in hiPSC-CM cultured with different concentrations of RA (concentration given on top). Gray lines indicate individual cells (number of cells given in brackets, dissociated from 3 EHTs). Open circles indicate mean values±SD. * indicates p<0.05 repeated measures ANOVA vs. respective basal values, # p<0.05 one-way ANOVA of log transformed data.

### Large transient outward currents are needed to recapitulate the high repolarization fraction and low plateau voltage typical for human atrium

To investigate whether the RA-induced increase in outward currents is able to reproduce the typical spike and dome shape of the human atria, we measured AP by sharp microelectrodes in intact EHT casted from hiPSC-CM differentiated in the absence and in the presence of different concentrations of RA. EHTs from hiPSC-CM differentiated in the presence of RA beat faster without differences between RA groups (**Figure 2B**). Take-off potentials were less negative in EHT cultured with 0.1 and 1 µM RA than in those cultured with 0.01 µM or in the absence of RA (**Figure 2**). Effects of RA were more pronounced on action potential duration (APD). Even with low RA concentration (0.01 and 0.1 µM) APD_90_ was shorter than in the control group. Similar shortening of APD_20_ and APD_90_ resulted in an unchanged repolarization fraction (APD_90_-APD_20_/APD_90_) in these two groups (**Figure 2**). In contrast, EHT casted from hiPSC-CM differentiated in the presence of 1 µM RA had a drastically shorter APD_20_ without further shortening of APD_90_, leading to the typical spike and dome AP shape of adult human atrium and a lower repolarization fraction (**Figure 2**). Thus, a critical concentration of RA (1 µM) is necessary and sufficient to induce a typical human atria-like AP shape in hiPSC-aCM in the EHT format.

**Figure 2:**
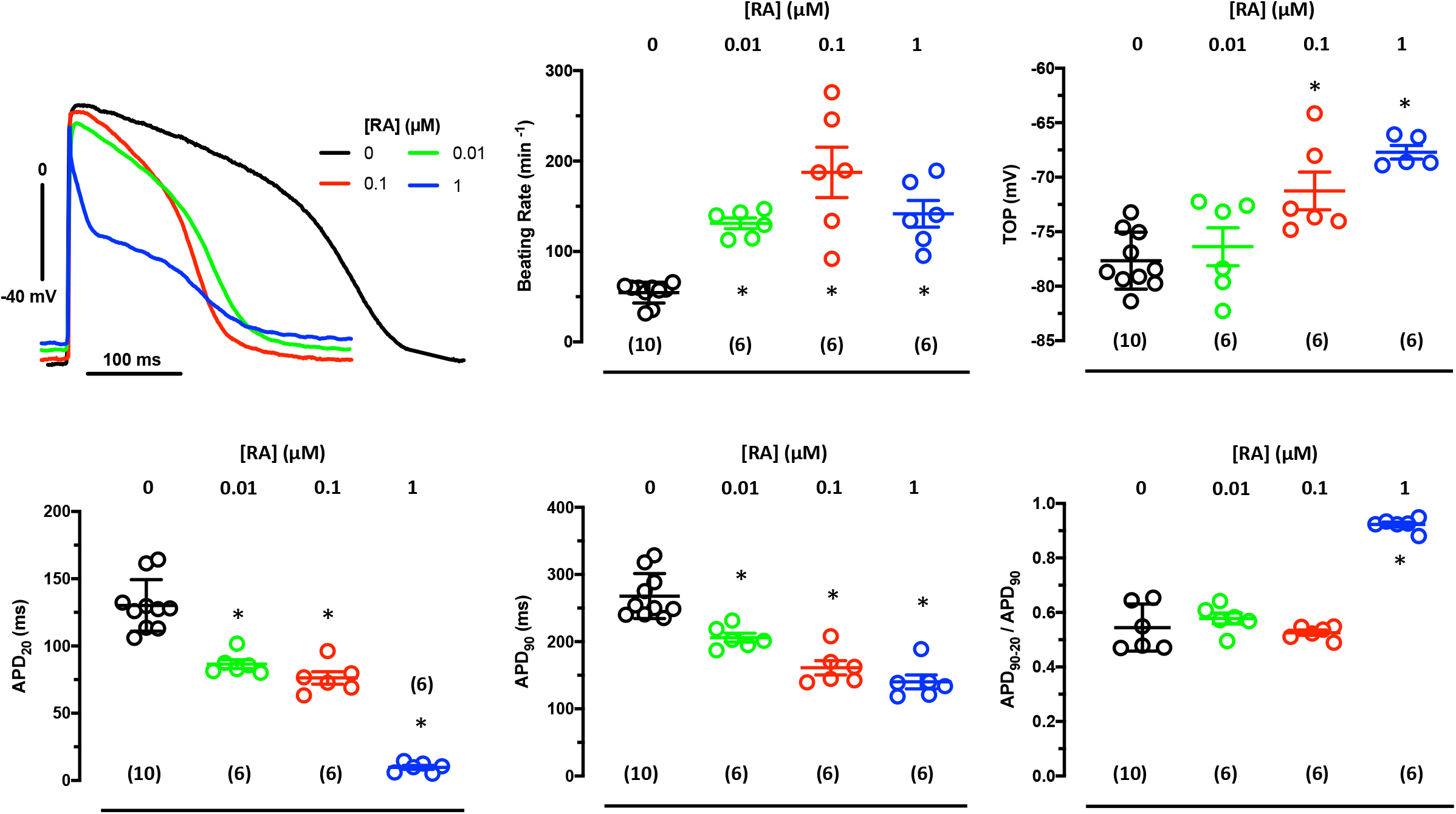
Concentration-dependency of RA on AP in EHT. **Top left**: Superimposed original traces of AP recorded in EHT based on hiPSC-CM cultured in the absence (0) or presence of RA concentrations. **Summary of results**: Mean values±SD for beating rate (**BR**), take-off potential (**TOP**), action potential duration (at 20 and 90% repolarization, **APD**_**20**_ and **APD**_**90**_) and repolarization fraction (**APD**_**90**_**-APD**_**50**_**/APD**_**90**_). * indicates p<0.05 vs. 0 RA, one way ANOVA of log transformed data. Number of EHTs resulting from one differentiation run is given in brackets.

### Large transient outward currents are required to recapitulate I_Kur_ block response pattern as seen in human atrium

In human atrium, block of I_Kur_ shifts the plateau voltage to less negative values and leads to a seemingly paradoxical abbreviation of terminal repolarization, i.e. decrease in APD_90_, because another important potassium current (I_Kr_) gets stronger activated at the less negative plateau voltage (Wettwer et al., 2004). To assess how much RA is needed to reproduce this pattern, we measured the effects of I_Kur_ block (50 µM 4-AP) in EHT casted from hiPSC-CM differentiated in the absence or presence of RA (0.01, 0.1 and 1 µM). In controls (0 RA), 50 µM 4-AP did not change APD (APD_20_ or APD_90_) or plateau voltage (**Figure 3**). In contrast, 4-AP prolonged APD_20_ in all three RA groups, indicating that even low concentrations of RA induce a relevant contribution of the atrial-selective I_Kur_ to repolarization. Only in EHT casted from hiPSC-CM differentiated in the presence of 1 µM RA, block of I_Kur_ shifted plateau voltage to less negative values accompanied by shortening in APD_90_, i.e. the canonical pattern of adult human atrium.

**Figure 3:**
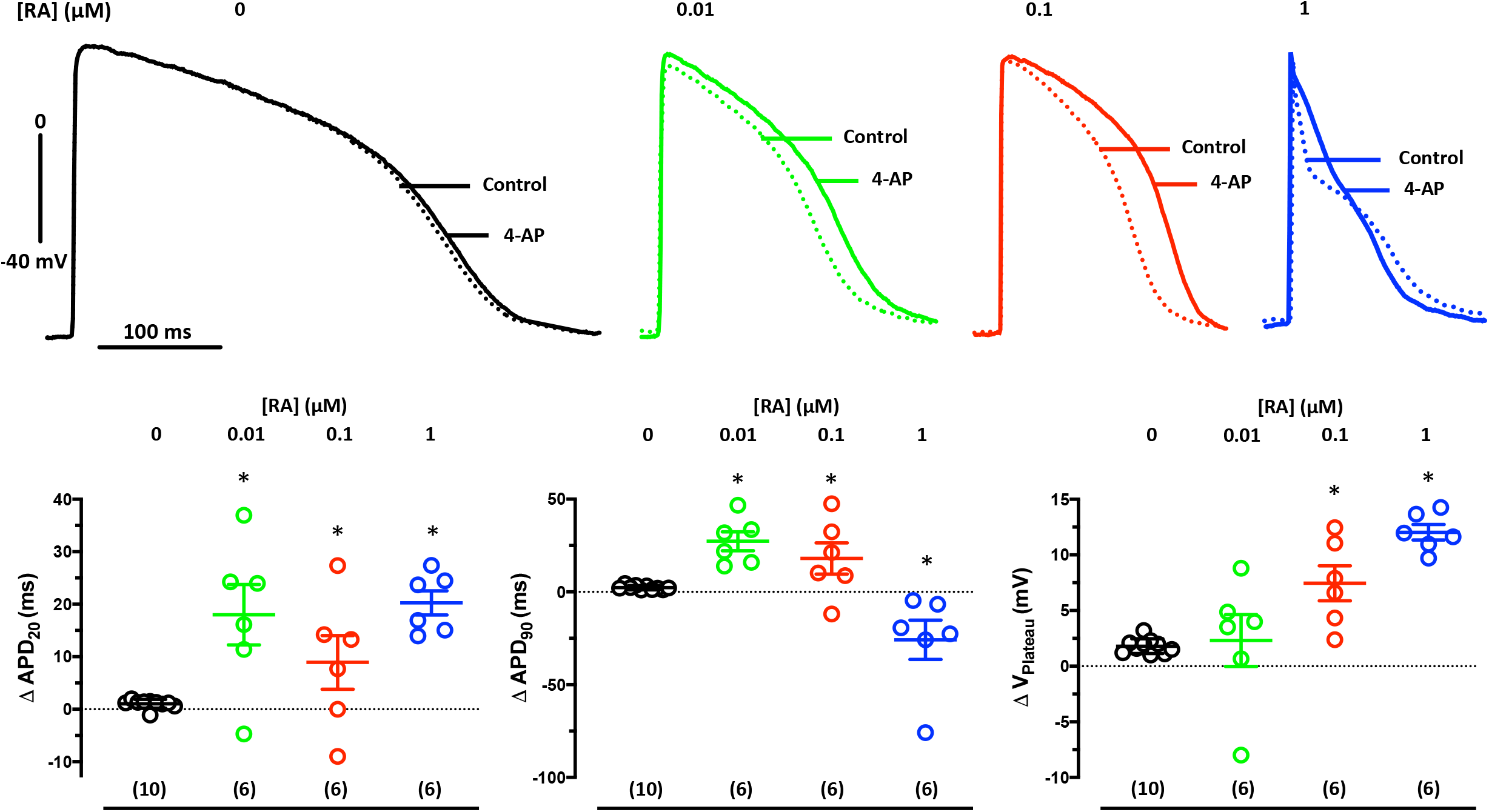
Concentration-dependency of RA on the AP-response to I_Kur_ block. **Top:** Superimposed original traces of AP before (Control) and after exposure to 50 µM 4-AP in EHT based on hiPSC-CM cultured in the absence (0 RA) or in the presence of RA (concentration given in brackets). **Bottom:** Summary of results for the effects of 4-AP on APD_20_ (**left**), on plateau voltage (V_Plateau_,**middle**) and on APD_90;_ (**right**). Open circles indicate mean values±SD. Numbers in brackets indicate number of EHTs resulting from one differentiation run. * indicates p<0.05 vs. 0 RA, one-way ANOVA of log transformed data.

To investigate whether shortening of APD_90_ upon I_Kur_ block in EHT (1 µM RA) is mediated by I_Kr_, we repeated experiments in the presence of the I_Kr_ blocker E-4031 (1 µM). I_Kr_ block alone prolonged APD_90_, but not APD_20_. E-4031 did not affect plateau voltage. As seen before in the absence of E-4031, I_Kur_ block by 50 µM 4-AP shifted plateau voltage to less negative values when given on top of E-4031. However, under this condition, 4-AP no longer shortened, but prolonged APD_90_ (**Figure 4**).

**Figure 4:**
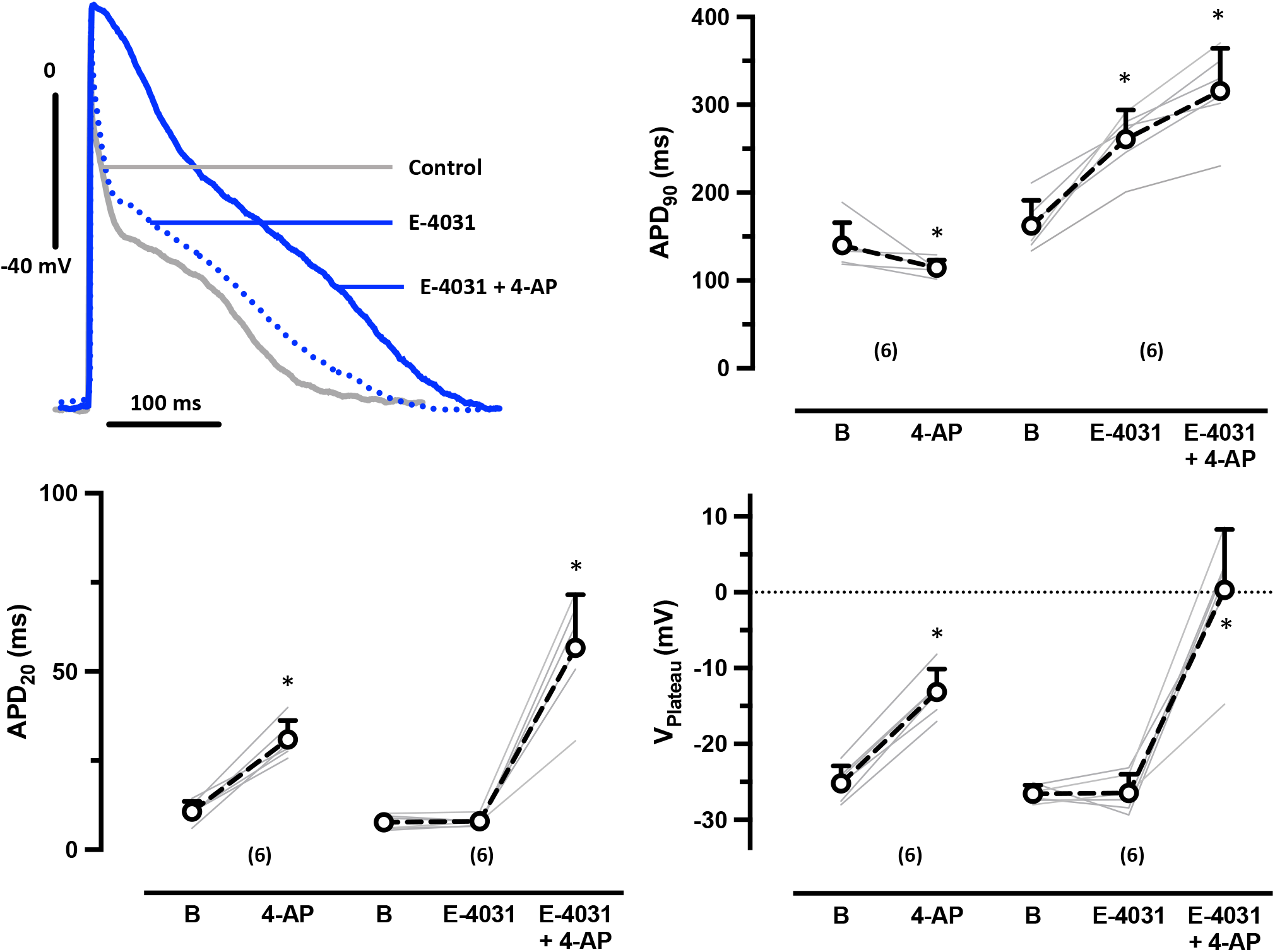
In aEHT (1 µM RA) the I_Kur_ block-induced shortening of APD_90_ depends on I_Kr_. **Top left:** Superimposed original traces of AP in EHT based on hiPSC-CM treated with 1 µM RA before (basal, B) and after exposure to 50 µM 4-AP in presence of the I_Kr_ blocker E-4031 (1 µM, right). Summary of results for the effects of 4-AP in the absence and presence of E-4031 on APD_90_ top, **left**), on APD_20_ (**bottom, left**) and on plateau voltage (V_Plateau_, bottom, **right**). Gray lines indicate individual EHT. Numbers in brackets indicate number of EHTs resulting from one differentiation run. Open circles indicate mean values±SD. * indicates p<0.05 vs. basal or E-4031, respectively (paired t-test).

### Low concentrations of RA are insufficient to produce relevant APD shortening upon muscarinic receptor activation

Besides I_Kur_, the physiology of human atria is characterized by the expression of large acetylcholine-sensitive inward rectifying ion currents, I_KACh_. To assess I_K,ACh_ contribution to the AP shape, we measured AP responses to 10 µM carbachol (CCh) in EHT from the three RA groups. We found a slight, but significant decrease in beating rate upon CCh in all three groups (**Figure 4**). However, only in EHT of the 1 µM RA group, CCh significantly decreased APD_90_ and shifted take-off potential (TOP) to more negative values. To investigate whether 1 µM RA has an uniform effect on the occurrence of I_K,ACh_ in hiPSC-CM differentiated with RA, we measured CCh responses of inward rectifier currents in single hiPSC-aCM (isolated from EHT of the 1 µM RA group only). Current density, measured at -100 mV, before adding CCh was 7.7±0.9 pA/pF (n=34). We saw a rapid increase in inward rectifier current upon exposure to CCh **(Figure 6)**. There was a large scatter in effect size between individual cells, but any cells showed an increase. Mean current density was more than doubled with 18.9±2.1 pA/pF (p<0.05, paired t-test), indicating robust and uniform expression of I_K,Ach_ in hiPSC-CM when differentiated in the presence of 1 µM RA.

**Figure 5:**
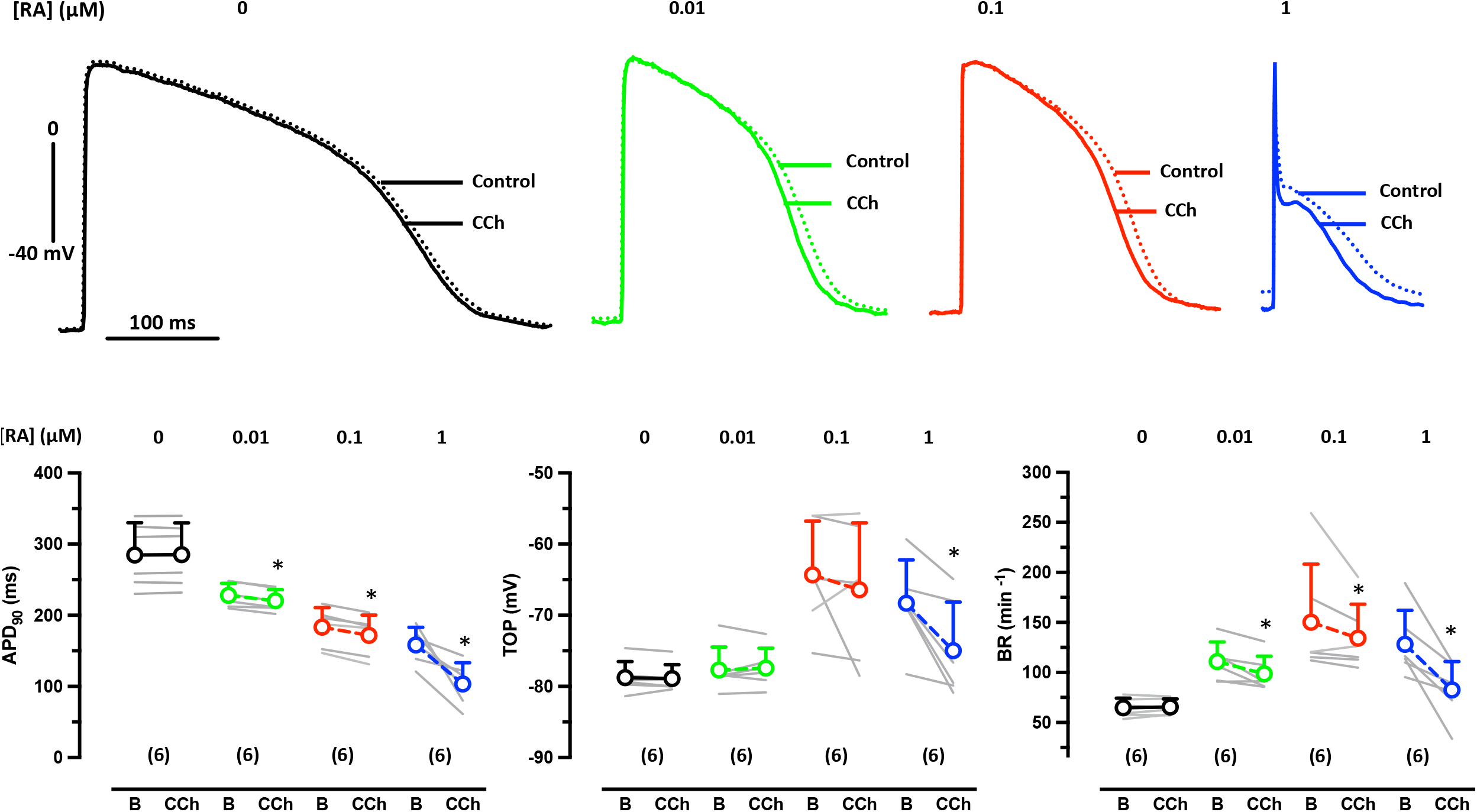
Concentration-dependency of RA on carbachol effects on APD_90_ in EHT. **Top**: Superimposed original traces of AP before (basal, B) and after exposure to 10 µM carbachol (CCh) recorded in EHT based on hiPSC-CM cultured in the presence of 0.01, 0.1 and 1 µM RA. **Bottom**: Summary of results for the effects of CCh on beating rate (BR, **left**), take-off potential (TOP, **middle**) and APD_90_ (**right**). Gray lines indicate individual EHT. Numbers in brackets indicate number of EHTs resulting from one differentiation run. Open circles indicate mean values±SD. * indicates p<0.05 vs. basal (paired t-test). Number of EHTs is given in brackets.

**Figure 6:**
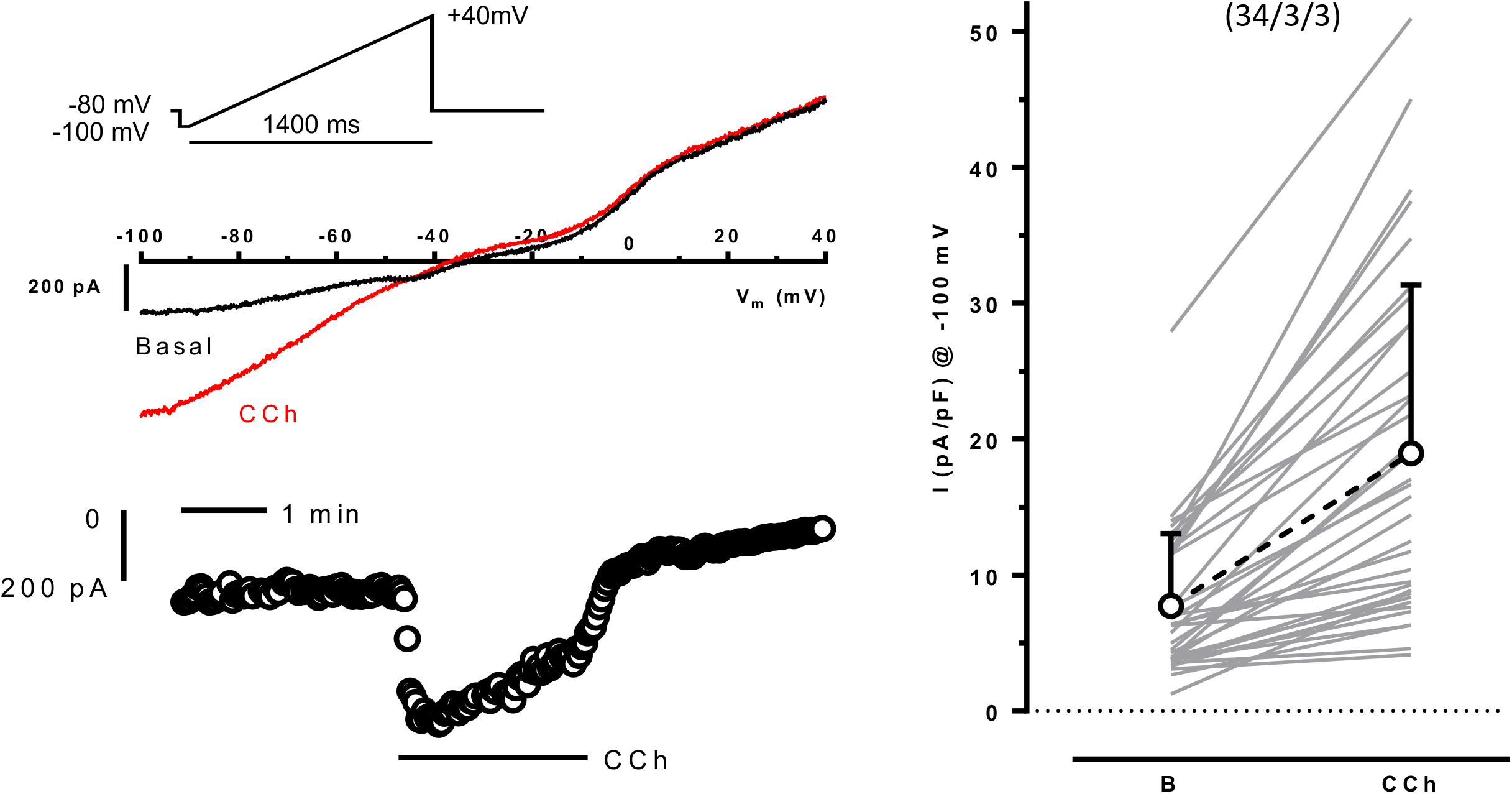
Consistent increase of inward rectifier currents upon carbachol in hiPSC-CM (1 µM RA) **Top left**: Superimposed original current traces evoked by a slow voltage ramp in a hiPSC-CM cultured in the presence of 1 µM RA before (Basal) and after exposure to 10 µM carbachol. Pulse protocol given as inset. **Bottom left:** Time course of the current at -100 mV in response to CCh. **Right:** Summary of results for the current at -100 mV. Gray lines indicate individual hiPSC-aCM, open circles indicate respective mean values±SD. Numbers in brackets indicate number of cells/number of dissociated EHTs, * indicates p<0.05 vs. basal (paired t-test).

**Figure 7:**
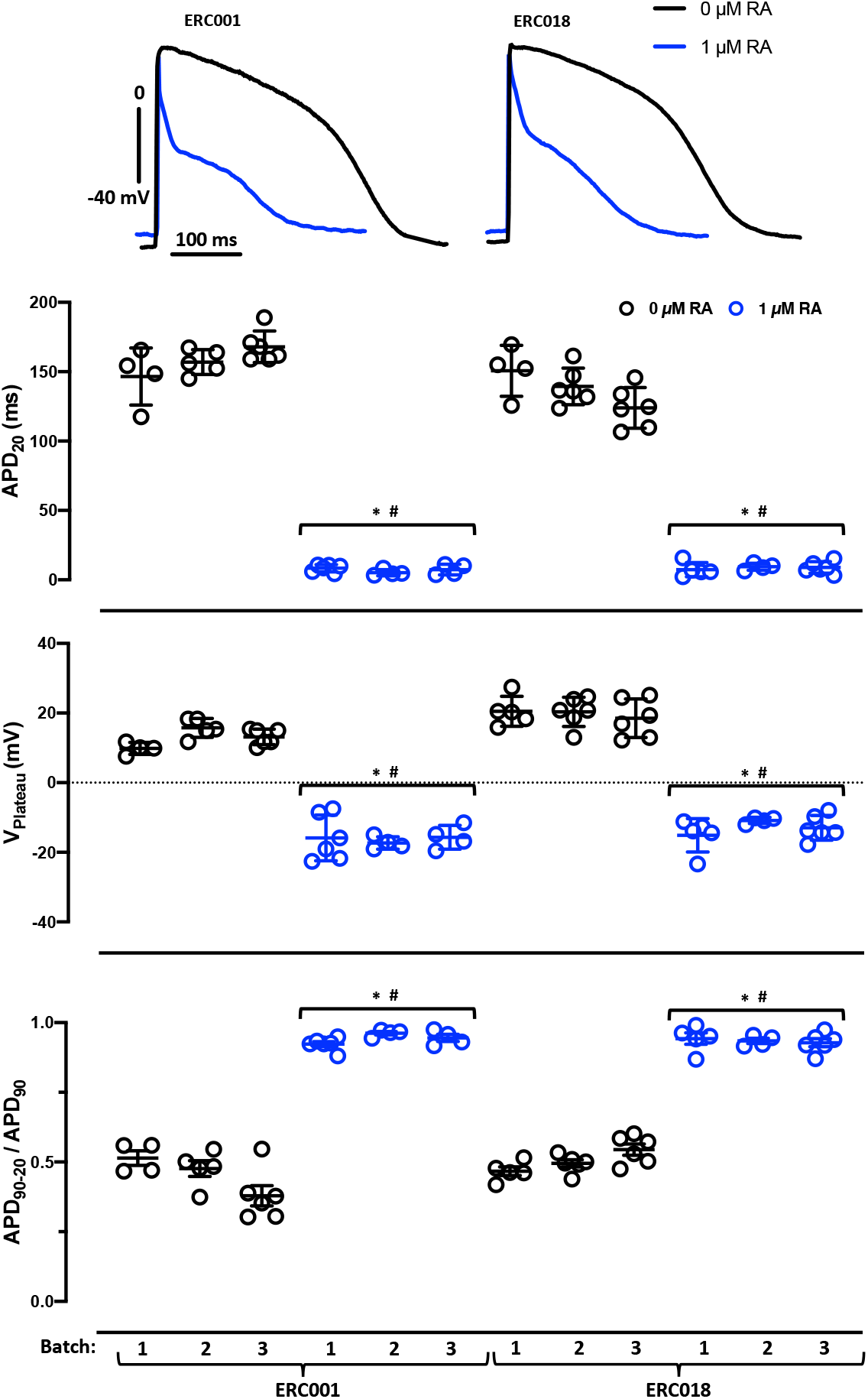
Reproducibility of RA-effects on AP shape. Top: Original recordings of AP recorded in EHT based on hiPSC-CM treated with 1 µM RA or not from two different cell lines (ERC001 and ERC018). Bottom: Individual data points and mean values±SD for APD_20,_ plateau voltage (V_Plateau_,) and repolarization fraction (APD_90_-APD_20_/APD_90_,**D**) in EHT from three different batches of RA-treated (1 µM RA) vs. untreated hiPSC-CM (0 RA) from two different cell lines. * indicates p<0.05 for nested ANOVA vs. 0 RA of the same cell line. # indicates no significant difference between single batches (ANOVA, every batch against another).

### Loss of RA by sterile filtration

Given the obvious discrepancies between the present results and our own study using a very similar methodology (Lemme et al. 2018), we carefully compared experimental procedures and identified sterile filtration of stock solution of RA in the prior study to be the only apparent difference. We suspected sterile filtration as a critical step leading to substantial loss of RA concentration in stock solutions. We therefore measured effect of sterile filtration on RA concentration by three different filters. All three filters adsorbed a significant amount of RA (**Table 1**). When using a PETF filter RA recovery was only about 0.1%.

**Table 1:** Loss of RA by sterile filtration. Mean values±SD for recovery rate of RA concentration in % of unfiltrated controls (n=6, data not shown) for three different filters.

|  | Filtropur S® (n=9) | Millex-GP® (n=9) | Whatman® REZIST (n=9) |
| --- | --- | --- | --- |
| Recovery in % | 29.8±2.4 | 28.6±7.6 | 0.1±0.01 |

### Reproducibility of RA-effects on AP shape

We have addressed the issue of reproducibility of RA-treatment on AP shape in a larger number of EHT prepared from three different batches (both RA treated and non-treated). In addition, experiments were done in as second, independent cell line (ERC018). RA-treatment induced an atrial-like AP shape (V_Plateau_ < 0 mV, APD_20_ < 15 ms and repolarization fraction close to 0) in all three batches of both cell lines. There was no overlap in the selected parameters between EHT based on hiPSC.CM treated or not treated with RA

## Discussion

Our study has two implications:

1. A concentration of 1 µM RA is sufficient and necessary to induce a fully human-like atrial AP phenotype in hiPSC-CM in the EHT format.
2. Sterile filtration of RA can lead to a relevant loss of RA loss, preventing induction of the adult-like atrial AP phenotype.

RA is crucial to induce atrial differentiation in hiPSC-CM, and several groups have used the shape of AP as proof of an atrial phenotype in RA-treated hiPSC-CM (**Table 2**). While it is clear that all studies show RA-induced effects on AP parameters towards a human atrial phenotype when compared to CM from standard differentiation protocols, the “atrial myocyte-likeness” varies widely. We will discuss some points that could be relevant for the inconsistencies.

**Table 2:** Action potential parameters in different types of atrial cardiomyocytes. Overview about action potential (AP) parameters of different cell types and different studies. The different cell types included adult atrial tissue, cardiomyocytes differentiated from human induced pluripotent stem cells (hiPSC-CM) and cardiomyocytes differentiated from human embryonic stem cells (hESC-CM). The effects of carbachol (CCh) and 4-aminopyridine (4-AP) are given as percentage of baseline values. * indicates an estimated parameter from AP shape.

| Cell type | Adult atrial tissue | Adult atrial tissue | Adult atrial tissue | hiPSC-CM | hiPSC-CM | hiPSC-CM | hiPSC-CM | hiPSC-CM | hESC-CM | hESC-CM | hESC-CM |
| --- | --- | --- | --- | --- | --- | --- | --- | --- | --- | --- | --- |
| Culture format |  |  |  | 3D | 3D | 3D | 3D | 2D | 2D | 2D | 2D |
| RA concentration ($\mu\text{M}$) | | | | 1 | 1 | 1 | 0.25-0.5 | 2 | 1 | 1 | 1 |
| Recording technique | Sharp ME | Sharp ME | Sharp ME | Sharp ME | Sharp ME | VD | Patch clamp | Patch clamp | Patch clamp | Patch clamp | Patch clamp |
| Temperature | 37 °C | 37 °C | 37 °C | 37 °C | 37 °C | RT | 31 °C | RT | 36 °C | RT | RT |
| MDP/RMP (mV) | -74 | -75 | -76 | -68 | -69 |  | -63 | -55 | -72 | -56 | -56 |
| APA (mV) | 95 |  |  | 82 |  |  |  | 85 | 80 | 82 | 80 |
| APD<sub>20</sub> (ms) | 7.2 | 5 | 8 | 10 | 31 |  | 115* | 13* | 21 | n. d. | 37* |
| APD<sub>90</sub> (ms) | 317 | 414 | 314 | 140 | 220 |  | 205 | 189 | 145 | 169 | 247 |
| APD<sub>20</sub>/APD<sub>90</sub> | 0.02 | 0.01 | 0.02 | 0.07 | 0.14 | 0.12* | 0.56* | 0.07 | 0.14 | n. d. | 0.14 |
| dV/dt<sub>max</sub> (V/s) | 219 |  |  | 104 |  |  | 7 | 68 | 26 | 13 | 63 |
| V<sub>Plateau</sub> | -16 | -21 |  | -16 |  |  |  |  | -10 |  |  |
| **4-AP effect** |  |  |  |  |  |  |  |  |  |  |  |
| APD<sub>20</sub> (% of baseline) |  | 240 | 194 | 300 | 106 |  |  |  | 205 |  |  |
| APD<sub>90</sub> (% of baseline) |  | 84 | 88 | 81 | 100 |  |  |  | 120 |  |  |
| V<sub>PLT</sub> (% of baseline) |  | 28 |  | 25 | 100 |  |  |  | 140 |  |  |
| **CCh effect** |  |  |  |  |  |  |  |  |  |  |  |
| APD<sub>90</sub> (% of baseline) |  |  | 55 | 60 | 82 |  | 69 |  | 97 |  |  |
| MDP/RMP (% of baseline) |  |  | 102 | 110 | 107 |  |  |  | 107 |  |  |
| Author/Year | Ravens 2015 | Wettwer 2004 | Lemme 2018 | This study | Lemme 2018 | Cyganek 2018 | Goldfracht 2020 | Lee 2017 | Devalla 2015 | Zhang 2011 | Laksman 2017 |

### Effects of cell source (hESC vs. hiPSC) and culture conditions (3D vs. 2D)

It seems reasonable to suspect an impact of cell source on the success of RA to induce an atrial AP phenotype. Three out of seven papers that reports in detail effects of RA on AP shape have used hESC (Devalla et al., 2015; Laksman et al., 2017; Zhang et al., 2011), the other four studies hiPSC (Cyganek et al., 2018; Goldfracht et al., 2020; Lee et al., 2017; Lemme et al., 2018). Data on APD_20_ are available from most studies (**Table 2**). APD_20_ was consistently short in both studies based on hESC (Devalla et al., 2015; Laksman et al., 2017), but varied widely when hiPSC were used. The data could indicate that RA effects are less robust in hiPSC, but the current data in hiPSC argue against this idea. Alternatively, the biology of individual hiPSC lines may vary more than that of hESC lines.

Direct head-to-head comparisons of 2D and 3D culture conditions are, to the best of our knowledge, limited to an earlier study of our group, but here the effects of RA were more pronounced in 3D EHTs than 2D monocultures, both in terms of atrial gene expression patterns and repolarization fraction (Lemme et al., 2018). The fact that repolarization fraction was higher in 3D than in 2D even in the absence of RA indicates that the 3D format itself contributes to improved atrial phenotype in 3D (Lemme et al., 2018). On the other hand, three out of four studies on RA-treated hESC-CM/hiPSC-CM cultured in 2D format reported APD_20_ values between 13 and 37 ms, close to situation in human atrium (Devalla et al., 2015; Laksman et al., 2017; Lee et al., 2017), indicating that RA is able to induce a strong, atrial-like early repolarization also in the 2D format.

### Recording techniques and other methodological conditions

We measured AP in intact EHT with sharp microelectrodes, while AP in hESC-aCM and hiPSC-aCM were mostly assessed by patch clamp recordings in isolated cells. This methodological difference may count. We observed previously that data scatter of APD and RMP was much larger when AP were measured in isolated cells compared to intact tissues (Horváth et al., 2018). The parameter repolarization fraction loses its power to discriminate between adult human ventricle and atrium when AP were measured in individual cells (Horváth et al., 2018).

In contrast, enzymatic isolation maybe an important reason for the observed differences. Devalla et al. (Devalla et al., 2015) reported a short APD_20_ and a strong effect of 4-AP at 50 µM (including an upwards shift of V_Plateau_ like in this study and in human adult atrium), but APD_90_ was prolonged in the presence of 4-AP instead of the expected shortening. This finding strikingly resembles the situation we report here obtained in the presence of E-4031. The I_Kur_ block-induced shortening of APD_90_ depends on I_Kr_, both in human adult atrium (Wettwer et al., 2004) and in hiPSC-aCM (**Figure 4**). Importantly, the hERG channel that mediates I_Kr_ can be destroyed by enzymes frequently used for cell isolation. In fact, I_Kr_ currents disappeared within minutes when cells were superfused with protease XIV, protease XXIV, proteinase K, or trypsin (Rajamani et al., 2006). Devalla et al. used a commercial kit (TrypLE™ Select Enzym, catalogue no. 12563011, ThermoFisher, Waltham, Massachusetts, USA) that contains a not further defined protease. Therefore, it seems justified to speculate that enzymatic dissociation of hiPSC-aCM may have abolished the contribution of I_Kr_ to AP regulation and thereby changed the response pattern to I_Kur_ block in their study.

### Loss of RA by sterile filtration

We saw a substantial loss of RA by sterile filtration. Effects are so large that we would no longer expect full effects of RA on differentiation. It is hard to determine if loss of RA by sterile filtration may explain the variability between different studies mentioned above since not only one study (including ours) has reported whether RA stock solutions or RA containing culture medium were sterile filtered or not. We felt safe to perform sterile filtration since the manufacturers states in its official product information that “Mahady and Beecher report sterile filtering RA solutions before addition to suspension cells” (Sigma, 1996). However, we were surprised that RA recovery after sterile filtration was not reported in the cited paper (Mahady and Beecher, 1996). No details on the filter material were given in that study. At least contemporarily used sterile filters absorb huge amounts of RA. Thus it seem wise to check recovery of RA or to refrain from sterile filtration of RA containing solutions.

## Limitations

We cannot exclude that changes in other currents like outward currents may have contributed to the effects of RA on repolarization. We haven’t used RA concentrations higher than 1 µM. Thus, it remains open whether higher RA concentration than 1 µM can have detrimental effects on AP shape in hiPSC-CM. We did not measure effects of RA treatment on gene expression.

## Experimental procedures

### Atrial differentiation of hiPSC and generation of atrial EHT

For all experiments the healthy in house control cell line ERC001 and ERC018 were used (UKEi001-A, https://hpscreg.eu/cell-line/UKEi001-A; UKEi003, https://hpscreg.eu/cell-line/UKEi003-C). All experimental methods for these procedures were approved by the Ethical Committee of the University Medical Center Hamburg-Eppendorf (Az. PV4798, 28.10.2014). All patients gave written informed consent. Protocols for hiPSC expansion, atrial cardiomyocyte differentiation and EHT generation for both hiPSC lines were performed as previously described (Breckwoldt et al., 2017). The hiPSC lines were derived from dermal fibroblasts from two healthy donors. In brief, embryoid bodies (EBs) were generated from expanded hiPSCs using spinner flasks and stirred suspension. Mesodermal induction was performed by growth factor cocktail (BMP-4 10 ng/ml, activin A 3 ng/ml, bFGF 5 ng/ml) for three days and the cardiac differentiation by WNT signal inhibitor XAV939 (1 μM). For the induction of atrial cardiomyocyte differentiation, RA (0.01, 0.1 or1 µM) was added for the first 72 hours of WNT signalling inhibition as recently described (Lemme et al., 2018). RA (Sigma Aldrich R2625) was dissolved in DMSO (Dimethyl Sulfoxide; stock concentration 100 μM) and used **without sterile filtration**. The differentiation run for all hiPSC treated with different RA-concentrations and for the respective control (0 RA) were prospectively performed in parallel.

After successful differentiation, dissociation of EBs was performed with collagenase II (200 U/L, Worthington, LS004176 in Hank’s balanced salt solution minus Ca^2+^/ Mg_2+_, Gibco, 14175-053 3.5 h, normoxia, 37 °C) (Breckwoldt et al., 2017). EBs were incubated with collagenase for 3.5 h (37 °C, normoxia) and were dispersed to isolated atrial cardiomyocytes (hiPSC-aCM).

Atrial-like engineered heart tissue (aEHT) was generated from 1 million hiPSC-aCM per construct. The fibrin gel matrix was made by mixing hiPSC-aCM, fibrinogen (Sigma F4753) and thrombin (100 U/L, Sigma Aldrich T7513) which were poured into agarose (1%) casting molds with silicone posts inserted from above (Breckwoldt et al., 2017; Hansen et al., 2010; Lemme et al., 2018).

### Action potential measurement

AP measurements were performed with standard sharp microelectrode as described previously.(Lemoine et al., 2018; Wettwer et al., 2013) All measurements were done in aEHTs which were continuously superfused with Tyrode’s solution (NaCl 127 mM, KCl 5.4 mM, MgCl_2_ 1.05 mM, CaCl_2_ 1.8 mM, glucose 10 mM, NaHCO_3_ 22 mM, NaHPO_4_ 0.42 mM, balanced with O_2_-CO_2_ [95:5] at 36°C, pH 7.4). The sharp microelectrode consisted of filamented borosilicate glass capillaries with an external diameter of 1.5 mm and internal diameter of 0.87 mm (HILG1103227; Hilgenberg, Malsfeld, Germany). The DPZ-Universal puller (Zeitz Instruments, Munich, Germany) was used to fabricate microelectrodes which had a resistance between 25 - 55 MΩ when filled with 2 M KCl. The pipettes were controlled by a hydraulic micromanipulator (Narishige MO-203) ensuring a delicate contact to the tissue. The aEHTs were transferred from the 24-well EHT culture plate into the AP measuring chamber by cutting the silicone posts and were fixed with needles in an optimal position for AP recording. The signals were amplified by a BA-1s npi amplifier (npi electronic GmbH, Tamm, Germany). APs were recorded and analysed using the Lab-Chart software (version 5, AD Instruments Pty Ltd., Castle Hill NSW, Australia). Definition of V_PLATEAU_ (Ford et al., 2013) was slightly modified as the voltage at in the range of ±5 ms time around 30% of APD_90_. Take-of potential (TOP) was defined as the diastolic membrane potential directly before the upstroke.

### Current measurements

Ion currents were measured at 37 °C using the whole-cell configuration of the patch-clamp technique by an Axopatch 200B amplifier (Axon Instruments, Foster City, CA, USA). The ISO2 software was used for data acquisition and analysis (MFK, Niedernhausen, Germany). Heat-polished pipettes were pulled from borosilicate filamented glass (Hilgenberg, Malsfeld, Germany). Tip resistances were 2.5– 5 MΩ, seal resistances were 3–6 GΩ. Cell capacitance (C_m_) was calculated from steady-state current during depolarizing ramp pulses (1V/1s) from −40 to −35 mV. Human iPSC-CMs used for patch clamp measurements were dissociated from EHT by collagenase II (200 U/mL; Worthington Biochemical, Lakewood, NJ, USA) for 5 h. Isolated cells were plated on gelatine-coated (0.1%) glass coverslips (12 mm diameter; Carl Roth GmbH + Co, Karlsruhe, Germany) and kept in culture for 24– 48 h to maintain adherence under superfusion in the recording chamber during patch clamp measurements. The cells were investigated in a small perfusion chamber placed on the stage of an inverse microscope. Inward rectifier currents were measured with the following bath solution (in mM): NaCl 120, KCl 20, HEPES 10, CaCl_2_ 2, MgCl_2_ 1 and glucose 10 (pH 7.4, adjusted with NaOH. Contaminating Ca^2+^ currents were suppressed with the selective L-type calcium channel blocker nifedipine (10 µM). The internal solution included (in mM): DL-Aspartate potassium salt 80, KCl 40, NaCl 8, HEPES 10, Mg-ATP 5, Tris-GTP 0.1, EGTA 5 and CaCl_2_ 2, pH 7.4, adjusted with KOH. Inward Current amplitudes were determined as currents at –100 mV(Horváth et al., 2018). A single concentration (2 µM) of the muscarinic receptor agonist carbachol (CCh) was used to evoke I_K,ACh_. Transient outward currents were measured in a slightly modified bath solution (KCl 5.4 instead of 20). Currents were elicited by 500 ms long test pulses to +50 mV applied every five seconds from a holding potential of -60 mV (Christ et al., 2008). Cells were exposed to two concentrations of 4-AP (50 µM and 5 mM).

### Sterile filtration and mass spectrometry

We prepared RA stock solutions (100 µM) in DMSO. One mL of this solution was filtrated through 3 different sterile filters: Filtropur S^®^ (pore size 0.2 µm, polyethersulfon), Saarstedt, Nümbrecht, Germany; Millex-GP^®^ (pore size 0.22 µM, polyethersulfon), Merck Millipore Ltd., Cork, Ireland and Whatman^®^ REZIST (pore size 0.2 µm, polytetrafluorethen), Cytiva, Marlborough, MA, USA. Unfiltrated solution was used as control.

RA was quantified by liquid chromatography-tandem mass spectrometry (LC-MS/MS) as described previously (Morgenstern et al., 2021). Stock solutions of retinoic acid and internal standard, i.e. all-trans retinoic acid-d5 (Cayman Chemical, Ann Arbor, MI, USA), were made up in DMSO, respectively (1 mg/mL). Calibration curves ranged from 10 ng/mL to 1000 ng/mL, five levels. 20 μL of calibrator or sample were added to 1.5 ml Eppendorf tubes and 20 μL of all-trans retinoic acid-d5 at 1000 ng/mL were added to each tube and vortexed briefly. 200 μL of acetonitrile was added to each tube and vortexed for 1 min prior to centrifugation for 10 minutes at 13000 rpm. 100 µL of supernatant was transferred to an MS 96well plate with 20 µL of water and capped. LC was performed applying a gradient of 0.8 mL/min 25/75%, vol/vol%, A (0.1 % formic acid) and B (acetonitrile/methanol, 50/50), to 2/98% over 2:50 min:sec on a Luna™ 5 µm C18 50x2.0 mm 100 A column (Phenomenex, Aschaffenburg, Germany). For MS/MS analyses in the positive electrospray ionisation mode on a Varian 1200 TSQ (Agilent Technologies, Santa Clara, CA, USA) the transitions m/z 301.2>123.0 @18 eV and m/z 306.2>127.0 @18 eV were monitored for retinoic acid and internal standard, respectively and concentrations were calculated by peak area ratio determination of calibrators and samples.

### Statistics

Statistical analyses were performed by using GraphPad Prism software version 7 (GraphPad Software, San Diego, CA, USA). Data are presented as mean ± SD. Log-transformation was used to allow application of parametric testing of data (Ismaili et al., 2020). Statistical significance was considered for differences with a value of p < 0.05.

## Author contribution

C.S., J.K., B.P., A.H., T.E., D.A. and T.C. planned experiments. C.S., M.S. T.E., D.A. and T.C. contributed to experiments and data analysis. C.S. T.E. and T.C. wrote the manuscript. All authors approved the final version of the manuscript.

## Acknowledgments

Authors thank Anna Steenpaß for excellent help with patch clamp measurements and technical support in action potential recordings. We are grateful to Birgit Klampe and Thomas Schulze for help with CM differentiation.

## References

Amos, G.J., Wettwer, E., Metzger, F., Li, Q., Himmel, H.M., and Ravens, U. (1996). Differences between outward currents of human atrial and subepicardial ventricular myocytes. J. Physiol. 491, 31– 50.

Argenziano, M., Lambers, E., Hong, L., Sridhar, A., Zhang, M., Chalazan, B., Menon, A., Savio-Galimberti, E., Wu, J.C., Rehman, J., et al. (2018). Electrophysiologic Characterization of Calcium Handling in Human Induced Pluripotent Stem Cell-Derived Atrial Cardiomyocytes. Stem Cell Reports 10, 1867–1878.

Breckwoldt, K., Letuffe-Brenière, D., Mannhardt, I., Schulze, T., Ulmer, B., Werner, T., Benzin, A., Klampe, B., Reinsch, M.C., Laufer, S., et al. (2017). Differentiation of cardiomyocytes and generation of human engineered heart tissue. Nat. Protoc. 12, 1177–1197.

Burashnikov, A., Mannava, S., and Antzelevitch, C. (2004). Transmembrane action potential heterogeneity in the canine isolated arterially perfused right atrium: Effect of IKr and I Kur/Ito block. Am. J. Physiol. - Hear. Circ. Physiol. 286, 2393–2400.

Christ, T., Wettwer, E., Voigt, N., Hála, O., Radicke, S., Matschke, K., Várro, A., Dobrev, D., and Ravens, U. (2008). Pathology-specific effects of the I <inf>Kur</inf>/I <inf>to</inf>/I <inf>K,ACh</inf> blocker AVE0118 on ion channels in human chronic atrial fibrillation. Br. J. Pharmacol. 154.

Cyganek, L., Tiburcy, M., Sekeres, K., Gerstenberg, K., Bohnenberger, H., Lenz, C., Henze, S., Stauske, M., Salinas, G., Zimmermann, W.H., et al. (2018). Deep phenotyping of human induced pluripotent stem cell-derived atrial and ventricular cardiomyocytes. JCI Insight 3.

Devalla, H.D., Schwach, V., Ford, J.W., Milnes, J.T., El-Haou, S., Jackson, C., Gkatzis, K., Elliott, D.A., Chuva de Sousa Lopes, S.M., Mummery, C.L., et al. (2015). Atrial-like cardiomyocytes from human pluripotent stem cells are a robust preclinical model for assessing atrial-selective pharmacology. EMBO Mol. Med. 7, 394–410.

Ford, J., Milnes, J., Wettwer, E., Christ, T., Rogers, M., Sutton, K., Madge, D., Virag, L., Jost, N., Horvath, Z., et al. (2013). Human electrophysiological and pharmacological properties of XEN-D0101: A novel atrial-selective Kv1.5/IKur inhibitor. J. Cardiovasc. Pharmacol. 61, 408–415.

Goldfracht, I., Protze, S., Shiti, A., Setter, N., Gruber, A., Shaheen, N., Nartiss, Y., Keller, G., and Gepstein, L. (2020). Generating ring-shaped engineered heart tissues from ventricular and atrial human pluripotent stem cell-derived cardiomyocytes. Nat. Commun. 11, 1–15.

Gunawan, M.G., Sangha, S.S., Shafaattalab, S., Lin, E., Heims-Waldron, D.A., Bezzerides, V.J., Laksman, Z., and Tibbits, G.F. (2021). Drug screening platform using human induced pluripotent stem cell-derived atrial cardiomyocytes and optical mapping. Stem Cells Transl. Med. 10, 68–82.

Hansen, A., Eder, A., Bonstrup, M., Flato, M., Mewe, M., Schaaf, S., Aksehirlioglu, B., Schworer, A., Uebeler, J., Eschenhagen, T., et al. (2010). Development of a Drug Screening Platform Based on Engineered Heart Tissue. Circ. Res.

Heidbüchel, H., Vereecke, J., and Carmeliet, E. (1987). The electrophysiological effects of acetylcholine in single human atrial cells. J. Mol. Cell. Cardiol. 19, 1207–1219.

Honda, Y., Li, J., Hino, A., Tsujimoto, S., and Lee, J.K. (2021). High-Throughput Drug Screening System Based on Human Induced Pluripotent Stem Cell-Derived Atrial Myocytes ∼ A Novel Platform to Detect Cardiac Toxicity for Atrial Arrhythmias. Front. Pharmacol. 12, 1–12.

Horváth, A., Lemoine, M.D., Löser, A., Mannhardt, I., Flenner, F., Uzun, A.U., Neuber, C., Breckwoldt, K., Hansen, A., Girdauskas, E., et al. (2018). Low Resting Membrane Potential and Low Inward Rectifier Potassium Currents Are Not Inherent Features of hiPSC-Derived Cardiomyocytes. Stem Cell Reports 10, 822–833.

Ismaili, D., Geelhoed, B., and Christ, T. (2020). Ca2+ currents in cardiomyocytes: How to improve interpretation of patch clamp data? Prog. Biophys. Mol. Biol. 157, 33–39.

Laksman, Z., Wauchop, M., Lin, E., Protze, S., Lee, J., Yang, W., Izaddoustdar, F., Shafaattalab, S., Gepstein, L., Tibbits, G.F., et al. (2017). Modeling Atrial Fibrillation using Human Embryonic Stem Cell-Derived Atrial Tissue. Sci. Rep. 7, 1–11.

Lee, J.H., Protze, S.I., Laksman, Z., Backx, P.H., and Keller, G.M. (2017). Human Pluripotent Stem Cell-Derived Atrial and Ventricular Cardiomyocytes Develop from Distinct Mesoderm Populations. Cell Stem Cell 21, 179-194.e4.

Lemme, M., Ulmer, B.M., Lemoine, M.D., Zech, A.T.L.L., Flenner, F., Ravens, U., Reichenspurner, H., Rol-Garcia, M., Smith, G., Hansen, A., et al. (2018). Atrial-like Engineered Heart Tissue: An In Vitro Model of the Human Atrium. Stem Cell Reports 11, 1378–1390.

Lemoine, M.D., Krause, T., Koivumäki, J.T., Prondzynski, M., Schulze, M.L., Girdauskas, E., Willems, S., Hansen, A., Eschenhagen, T., and Christ, T. (2018). Human Induced Pluripotent Stem Cell-Derived Engineered Heart Tissue as a Sensitive Test System for QT Prolongation and Arrhythmic Triggers. Circ. Arrhythmia Electrophysiol. 11.

Mahady, G.B., and Beecher, C.W.W. (1996). Induction of benzo[c]phenanthridine alkaloid biosynthesis in suspension cell cultures of Sanguinaria canadensis by retinoic acid derivatives. Nat. Prod. Lett. 8, 173–180.

Morgenstern, J., Fleming, T., Kliemank, E., Brune, M., Nawroth, P., and Fischer, A. (2021). Quantification of all-trans retinoic acid by liquid chromatography–tandem mass spectrometry and association with lipid profile in patients with type 2 diabetes. Metabolites 11, 1–10.

Pei, F., Jiang, J., Bai, S., Cao, H., Tian, L., Zhao, Y., Yang, C., Dong, H., and Ma, Y. (2017). Chemical-defined and albumin-free generation of human atrial and ventricular myocytes from human pluripotent stem cells. Stem Cell Res. 19, 94–103.

Rajamani, S., Anderson, C.L., Valdivia, C.R., Eckhardt, L.L., Foell, J.D., Robertson, G.A., Kamp, T.J., Makielski, J.C., Anson, B.D., and January, C.T. (2006). Specific serine proteases selectively damage KCNH2 (hERG1) potassium channels and IKr. Am. J. Physiol. - Hear. Circ. Physiol. 290, 1278–1288.

Ravens, U., and Wettwer, E. (2011). Ultra-rapid delayed rectifier channels: Molecular basis and therapeutic implications. Cardiovasc. Res. 89, 776–785.

Sigma (1996). all trans-RETINOIC ACID Sigma Prod . No . R 2625.

Soepriatna, A.H., Kim, T.Y., Daley, M.C., Song, E., Choi, B.R., and Coulombe, K.L.K. (2021). Human Atrial Cardiac Microtissues for Chamber-Specific Arrhythmic Risk Assessment. Cell. Mol. Bioeng. 14, 441–457.

Wettwer, E., Hála, O., Christ, T., Heubach, J.F., Dobrev, D., Knaut, M., Varró, A., and Ravens, U. (2004). Role of IKur in controlling action potential shape and contractility in the human atrium: Influence of chronic atrial fibrillation. Circulation 110, 2299–2306.

Wettwer, E., Christ, T., Endig, S., Rozmaritsa, N., Matschke, K., Lynch, J.J., Pourrier, M., Gibson, J.K., Fedida, D., Knaut, M., et al. (2013). The new antiarrhythmic drug vernakalant: Ex vivo study of human atrial tissue from sinus rhythm and chronic atrial fibrillation. Cardiovasc. Res. 98, 145–154.

Wijffels, M.C.E.F., Kirchhof, C.J.H.J., Dorland, R., and Allessie, M.A. (1995). Atrial fibrillation begets atrial fibrillation: A study in awake chronically instrumented goats. Circulation 92, 1954–1968.

Zhang, Q., Jiang, J., Han, P., Yuan, Q., Zhang, J., Zhang, X., Xu, Y., Cao, H., Meng, Q., Chen, L., et al. (2011). Direct differentiation of atrial and ventricular myocytes from human embryonic stem cells by alternating retinoid signals. Cell Res.

