## Supplemented Material for "A critical role of retinoic acid concentration for the induction of a fully human-like atrial phenotype in hiPSC-CM"

| [RA] (µM) | BR | MDP | TOP | $APD_{20}$ | $APD_{90}$ | $APD_{90-20}$ $/APD_{90}$ |
|---|---|---|---|---|---|---|
| **0** | 54±11.5 | -78.1±0.9 | -77.7±2.6 | 130.2±19.2 | 267.9±33.6 | 0.54±0.1 |
| **0.01** | 131.1±14.7 * | *-76.6±3.8* | -76.4±4.3 | 86.4±7.9 * | 206±16 * | 0.58±0.1 |
| **0.1** | 187.5±68.7 # | *-71.8±4.9* | -71.3±4.2 # | 76.3±11.4 | 161.2±26 # | 0.53±0 |
| **1** | 147±36 § | *-68.6±0.7* § | -67.7±1.4 | 8.8±3.6 § | 140.2±25.5 | 0.92±0 § |

**Supplement Table 1: Effects of different concentrations of retinoic acid (RA) on AP parameters.** Mean values±SD for beating rate (BR), maximum diastolic potential (MDP), take-off potential (TOP), action potential duration at 20% ($APD_{20}$) and 90% repolarization ($APD_{90}$) and repolarization fraction ($APD_{90-20}$ $/APD_{90}$) measured in intact EHT based on hiPSC-CM differentiated with different retinoic acid (RA) concentrations. * p<0.05 vs. "0" RA, # p<0.05 vs. 0.01 µM RA and § p<0.05 vs. 0.1 µM RA (ANOVA, same n numbers as in **Figure 2**).

| [RA] (µM) | $\Delta APD_{20}$ | $\Delta APD_{90}$ | $\Delta V_{Plateau}$ |
|---|---|---|---|
| **0** | 1.5±2 | 4.2±5.9 | 3.2±2.1 |
| **0.01** | 18±14.1 * | 27.3±12.4 * | 2.3±5.7 |
| **0.1** | 8.9±12.6 | 18.1±20.6 | 1.1±8.9 # |
| **1** | 20.26±5.6 | -25.8±25.9 § | 12±1.7 § |

**Supplement Table 2: Effects of different concentrations of retinoic acid (RA) on 4-amniopyridine effects.** Mean values±SD for the effects of 50 µM 4-aminopyridine (4-AP, expressed as Δ-values) on action potential duration at 20% repolarization ($APD_{20}$), plateau voltage ($V_{Plateau}$) and action potential duration at 90% repolarization ($APD_{90}$) measured in intact EHT based on hiPSC-CM differentiated with different retinoic acid (RA) concentrations. * p<0.05 vs. "0" RA, # ü<0.05 vs. 0.01 µM RA and § p<0.05 vs. 0.1 µM RA (ANOVA, same n numbers as in **Figure 3**).

| [RA] (µM) | $\Delta APD_{90}$ | ΔTOP | ΔBR |
|---|---|---|---|
| **0** | 3.1±4.2 | 0.1±2 | 3.3±4.3 |
| **0.01** | 4.3±8.4 * | -0.03±1.4 | -8.8±16 * |
| **0.1** | 0.1±9.3 | -3.6±5.4 | -15.9±25.3 |
| **1** | -57.1±33 § | -7.2±3.4 | -45.6±28.1 |

**Supplement Table 3: Effects of different concentrations of retinoic acid (RA) on carbachol effects.** Mean values±SD for the effects of 10 µM carbachol (CCh, expressed as Δ-values) on action potential duration at 20% repolarization ($APD_{20}$), take-off potential (TOP) and beating rate (BR) measured in intact EHT based on hiPSC-CM differentiated with different retinoic acid (RA) concentrations. * p<0.05 vs. "0" RA, # ü<0.05 vs. 0.01 µM RA and § p<0.05 vs. 0.1 µM RA (ANOVA, same n numbers as in **Figure 5**).
